# The universal mechanism of intermediate filament transport

**DOI:** 10.1101/251405

**Authors:** Amélie Robert, Peirun Tian, Stephen A. Adam, Robert D. Goldman, Vladimir I. Gelfand

## Abstract

Intermediate filaments (IFs) are a major component of the cytoskeleton that regulates a wide range of physiological properties in eukaryotic cells. In motile cells, the IF network has to adapt to constant changes of cell shape and tension. In this study, we used two cell lines that express vimentin and keratins 8/18 to study the dynamic behavior of these IFs. We demonstrated that both IF types undergo extensive transport along microtubules. This was an unexpected result as keratin filament remodeling has been described to depend on actin dynamics. We established the role of kinesin-1 in vimentin and keratin IF transport by knocking out KIF5B, the ubiquitous isoform of kinesin-1. Futhermore, we demonstrated that unlike typical membrane cargoes, transport of both types of IFs does not involve kinesin light chains, but requires the presence of the same region of the kinesin-1 tail, suggesting a unified mechanism of IF transport.

## INTRODUCTION

Depending on the tissue or cellular context, cells face different physiological and mechanical challenges that can be overcome by the cell- and tissue-specific expression of one or a combination of some of the 70 genes encoding intermediate filament proteins in human. For example mesenchymal cells typically express high levels of type III vimentin IF (VIF), the assembly and disassembly of which facilitate different aspects of cell migration (1-4). In contrast epithelial cells express a combination of type I and type II keratin IFs that are connected to desmosomes and hemidesmosomes to ensure the tight connection between cells in the epithelial sheet and between the cells and the basal membrane (5)

To accommodate constant changes of cell shape as cells contract, migrate or invade, IFs need to undergo profound and constant reorganization (reviewed in (6)). This reorganization is achieved by a combination of severing and re-annealing (7, 8) as well as intracellular translocation of mature IFs and their precursors. The first indication that IFs could be a cargo for microtubule-based motors came from the microinjection of pan-kinesin antibody that induced the retraction of the vimentin network (9). With the development of fluorescent probes and advanced live cell imaging techniques, several types of cytoplasmic IFs have been observed to move along microtubule tracks. Movement of neurofilaments has been observed in axons of cultured neurons (10). Our previous studies have shown that vimentin particles as well as mature filaments are transported along microtubules (8, 11, 12). Recently, GFAP and nestin IFs were also reported to move together with vimentin in migrating astrocytes (13). Among the 40 kinesins in mammals, the major microtubule motor kinesin-1 has been typically suggested to be involved in IF transport. Kinesin-1 in mammals is represented by three isoforms, KIF5A, KIF5B or KIF5C, with KIF5B being the most abundant ubiquitous version. Several reports suggested that various types of cytoplasmic IFs might be potential cargo for kinesin-1. In axons for example, neurofilaments are transported by KIF5A (14, 15), while in muscle, KIF5B has been reported to be essential for the delivery of desmin and nestin IFs to the growing tip of myotubes (16). Recently, knock down of *KIF5B* has been shown to reduce anterograde transport of IFs in migrating astrocytes (13).

Obviously absent from this list are the most abundant IFs present in epithelial cells, keratin IF. For these filaments, a different mode of transport based on actin dynamics has been proposed. In this model, keratin IFs undergo a constant cycle of assembly and disassembly that involves actin-dependent centripetal motion of keratin particles and filaments from the cell periphery, where filament particles are formed, to the perinuclear region. When filaments reach the perinuclear region, a fraction of keratin subunits are released and returned by diffusion to the cell periphery where another cycle of particle formation takes place (17-20). Although microtubule-dependent motion of keratin particles has been observed previously (21, 22), the contribution of microtubules and/or microtubule-based motors for keratin filament dynamics has been neglected as the rapid transport of fully polymerized keratin filaments has never been reported. In this work, we used a combination of photoconversion experiments and CRISPR/Cas9 genome editing of *KIF5B* to compare the dynamics of keratin and vimentin IFs and the role of microtubules and microtubule motors. Surprisingly, we found that the dynamic properties of both classes of IFs include transport of long mature filaments along microtubules by kinesin-1 and the same domain of the kinesin tail is involved in transport, strongly suggesting that all types of IFs move along microtubules using an identical mechanism.

## RESULTS

### Vimentin intermediate filaments are transported along microtubules by kinesin-1

Kinesin antibody injection and shRNA knock down experiments have suggested a role for kinesin-1 in vimentin IF transport (9, 13). In humans, kinesin-1 heavy chain is encoded by three genes; *KIF5A, KIF5B* and *KIF5C*. We used CRISPR/cas9 genome-editing to KO *KIF5B*, the major gene coding for kinesin-1 in RPE cells. Several clones were amplified and the KO was verified using western blot analysis with an antibody (CT) directed against a peptide in the tail domain of kinesin heavy chain common to all three isoforms of kinesin-1. Two of the clones were selected for further analysis. The specificity of the KO was further confirmed using a blot with an antibody (HD) that recognizes the motor domains of multiple kinesins. This blot demonstrated that only the band corresponding to kinesin-1 was absent from the lysates of the KO cells (Figure 1B). We checked the functional implications of *KIF5B* KO by analyzing the distribution and motility of known kinesin cargos. As expected, *KIF5B* KO induced the retraction of mitochondria from the cell periphery as described previously ((23), Figure S2 AB). In contrast, the motility of lysosomes was not affected (Figure S2C), as lysosome transport is driven, not only by kinesin-1, but by multiples kinesins (Reviewed in (24)).

**Figure 1.**
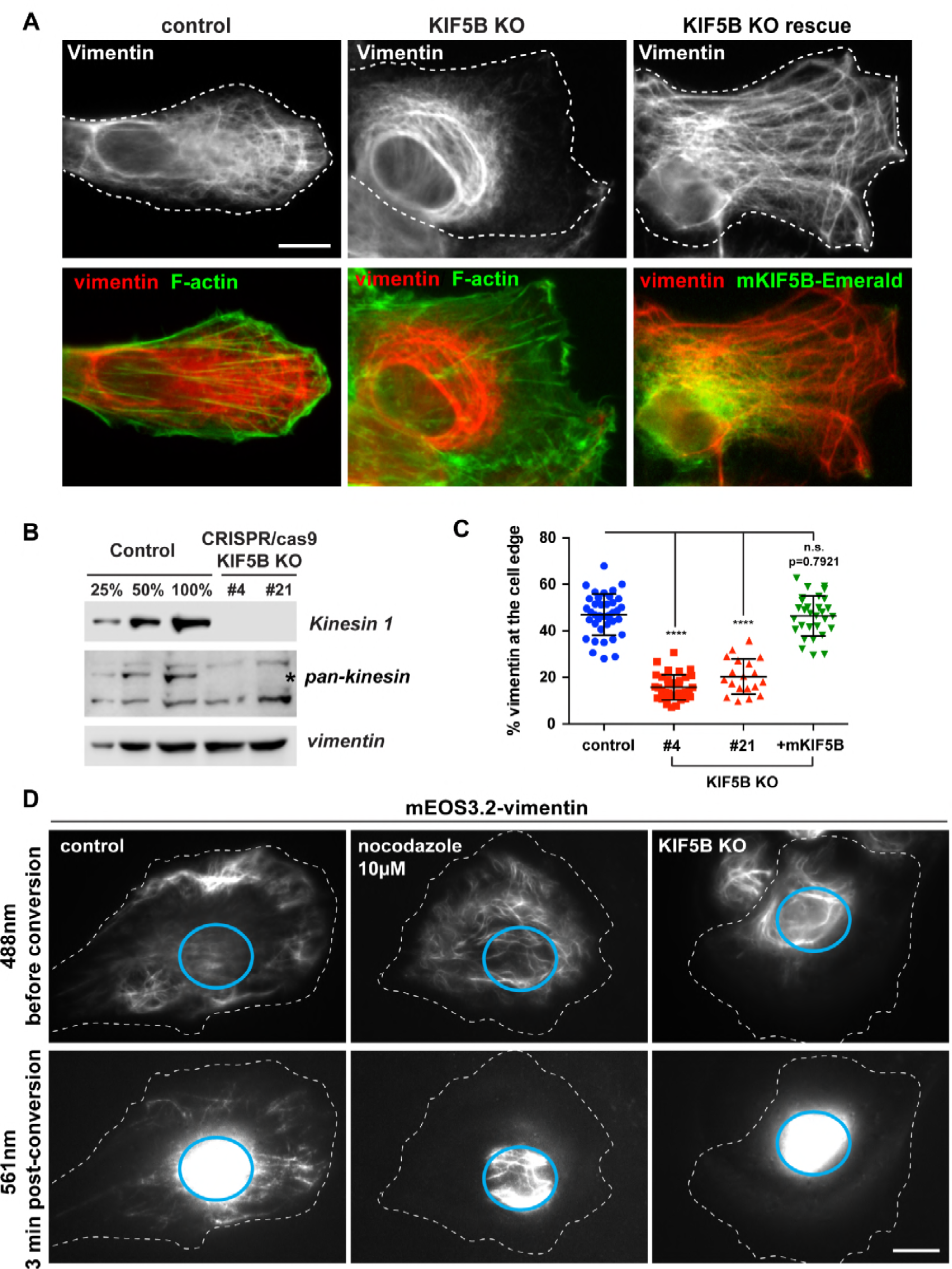
*KIF5B KO affects vimentin IF distribution and inhibits their transport A) Control and KIF5B KO cells were fixed and co-stained for vimentin and F-actin. The last column shows the rescue of vimentin distribution in a KIF5B KO cell by mKif5b-Emerald expression. B) Western blot analyses using kinesin-1 antibody shows the absence of kinesin-1 in two different KIF5B CRISPR clones. Pan-kinesin antibody shows the specificity of the KO for kinesin-1 (position marked by the star), but not the other kinesin family members. Vimentin is used as loading control. C) Graph shows the % of vimentin at the cell edge (Mean with SD; n >30 cells, except KIF5B KO #21 n=19). Data are representative of at least two independent experiments (see Materials and Methods for quantification details). Statistical significance was determined using the Mann-Whitney test (****; p<0.0001). D) Photoconversion of mEos3.2-vimentin in RPE cell. mEos3.2-vimentin was photoconverted from green to red at the cell center (cyan circle) and the dynamics of photoconverted filaments was imaged using TIRF microscopy. The top panels show the vimentin network in the green channel (488nm) before photoconversion and the bottom panels show the red channel (561nm) 3 minutes after photoconversion. Note the presence of several photoconverted filaments outside of the original photoconverted zone in the control cell; photoconverted filaments are confined inside the initial zone after microtubule depolymerization with nocodazole (10µM for 3 hrs) in KIF5B KO cells. Bars, 10µm.*

We performed vimentin immunostaining to determine how the absence of kinesin-1 impacts vimentin filament distribution. In control cells, the vimentin filament network extended all the way to the cell periphery as delineated by actin staining. In contrast, in the absence of KIF5B the majority of the mature vimentin filaments retracted from the leading edge, with only a few short IF and non-filamentous particles left behind (Figure 1A). We quantified the results using the procedure described in Figure S1, and confirmed the initial visual observation that the KO of *KIF5B* correlated with the retraction of the vimentin network towards the nucleus (Figure, 1C). To confirm that this phenotype was not due to an off-target effect, we performed a rescue experiment using a mouse version of KIF5B (*mKif5b*), which is insensitive to the gRNA used to KO human *KIF5B*. When mKif5b-Emerald was expressed in RPE *KIF5B* KO cells, the vimentin filament network distribution was fully restored, demonstrating that the retraction of the network was indeed caused by the absence of kinesin-1 (Figure 1A, third column). This result corroborates other observations (9, 11, 13) suggesting that vimentin is a kinesin-1 cargo.

We have previously visualized vimentin filaments transported along microtubules in RPE cells (8). To directly demonstrate that this transport is powered by kinesin-1, we used photoconversion of mEos3.2-vimentin (8). The emission of mEos3.2 changes from green to red when exposed to ultraviolet (UV) light at 400 nm. By restricting photoconversion to a circular area of about 10 μm in diameter, we produced fiduciary marks on filaments permitting us to monitor their transport in regions of cells with high filament density. Photoconverted mEos3.2-vimentin was imaged over a period of 3 min using TIRF microscopy. As described before (8), photoconverted filaments robustly moved away from the central photoconverted area (Figure 1D, left panel and Supplemental Video S1). When cells were treated with 10µM nocodazole for 3 hrs to depolymerize microtubules, vimentin filament transport was completely inhibited (Figure 1D, middle panels). To determine if the microtubule-dependent transport is driven by kinesin-1, mEos3.2 vimentin was expressed in RPE *KIF5B* KO cells and the same photoconversion experiment was performed. No converted filaments could be detected outside of the area of initial photoconvertion for 3 min (Figure 1D, right panels, Supplemental Video S2). This result is consistent with the effect of *KIF5B* KO on vimentin distribution, further demonstrating that kinesin-1 is the motor that drives transport of vimentin IFs along microtubules.

### Keratin intermediate filaments are transported along microtubules by kinesin-1

The role of the actin cytoskeleton in the cycle of keratin assembly/disassembly has been described in great details (Reviewed in (25)). However, even though microtubule-dependent motion of keratin particles has been observed previously (21, 22), the role of microtubules in the transport of keratin filaments has never been reported. In RPE cells, keratin filaments co-exist with vimentin filaments. This allows us to use our RPE KO cells for analysis of keratin transport. Immunostaining of the keratin network using a pan-keratin antibody shows an intricate network of keratin filaments that extend to the cell edge (Figure 2A). Interestingly, the absence of kinesin-1 causes the retraction of the keratin network from the cell periphery (Figure 2A-B). This observation suggests that at least in RPE cells, keratin filaments, like vimentin, can potentially be transported along microtubules by kinesin-1.

**Figure 2.**
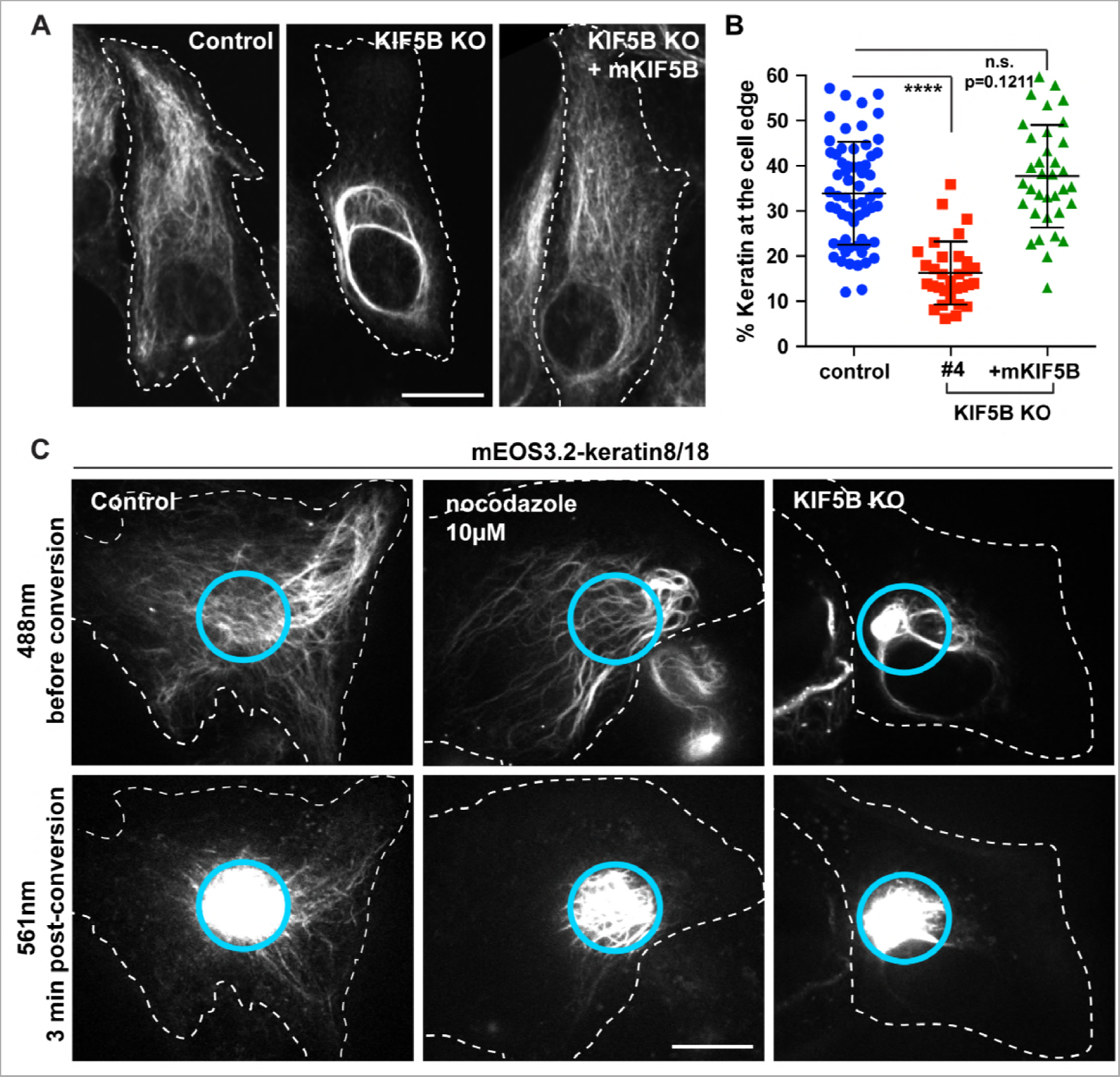
*KIF5B KO changes keratin filament distribution and inhibits keratin filament transport. A) Confocal imaging of keratin immunostaining in control versus KIF5B KO cells. The cell periphery was delineated by a dashed line to emphasize the retraction of the keratin filaments from the cell edge in KIF5b KO cells. In the last column, mKIF5B-Emerald was expressed in KIF5B KO cells to rescue keratin distribution. B) Graph shows the % of keratin at the cell edge (Mean with SD; n >30 cells). Data are representative of at least two independent experiments. Statistical significance was determined using the Mann-Whitney test (****; 159 p<0.0001). C) Photoconversion of mEos3.2-keratin 8/18 in RPE cells using spinning disk confocal microscopy. mEos3.2-keratin was photoconverted from green to red at the cell center (cyan circle). The top panels show keratin network in the green channel (488nm) before conversion and the bottom panels show the red channel (561nm) 3 minutes after photoconversion. Several photoconverted filaments were present outside of the original photoconverted zone in the control cells, while photoconverted filaments remained inside the initial zone after nocodazole treatement (10µM for 3hrs) in KIF5B KO cells. Bar, 10µm.*

To visualize keratin filament dynamics, we co-expressed photoconvertible keratin 8 and keratin 18 (mEos3.2-krt8/18) in RPE cells. As described for mEos3.2-vimentin, a subset of keratin filaments at the cell center was photoconverted and imaged in the red channel for 3 min using spinning disk confocal microscopy. Remarkably, long as well as short keratin filaments moved away from the photoconverted zone during 3 min (Figure 2C left panels, Video S3). The accumulation of keratin filaments outside the photoconverted zone was abolished by nocodazole treatment, demonstrating the role of microtubules in the transport of keratin filaments (Figure 2C, middle panels, Video S4). When the same experiment was conducted in *KIF5B* KO RPE cells, no transport of photoconverted keratin filaments outside of the photoconverted zone was observed (Figure 2C, right panels, Video S5). Therefore, kinesin-1 drives keratin filaments transport along microtubules.

### Keratin filaments are transported independently of vimentin

What distinguishes RPE from typical epithelial cells is that the keratin and vimentin IF networks look very similar, and often these two types of filaments are located in close proximity to each other (Figure S3A) raising the possibility that keratin filaments are co-transported with vimentin filaments. To test that hypothesis, we took advantage of CRISPR/Cas9 genome editing to KO vimentin in RPE cells (Figure S3D) and look at the impact of the absence of vimentin for keratin distribution. Surprisingly, in RPE cells the keratin filaments network completely collapsed in absence of vimentin, making it impossible to study keratin filament transport in vimentin null cells (Figure S3B). This phenotype is not caused by an off-target effect from the genome editing because it can be rescued by the expression of mouse vimentin insensitive to the gRNA used to KO human vimentin (Figure S3C-D).

Obviously, interdependence of two IF networks could be a unique feature of RPE cells because vimentin is absent from many keratin-expressing cells and thus is not universally required for normal keratin distribution. We, therefore, examined human carcinoma A549 cells, which co-express vimentin and keratin. In these cells, like in typical epithelial cells, the keratin IF network remained expanded even after vimentin KO (Figure 3A). First, we expressed mEos3.2-krt8/18 in A549 cells and observed that a subpopulation of keratin filaments was transported in these cells (Figure 3B, left panel, Video S6) showing that keratin transport along microtubules is not restricted to RPE cells but can occur in other cell types. Next a photoconversion experiment was performed in A549 vimentin KO cells. The result revealed that keratin filaments were transported even in the cells lacking vimentin (Figure 3B, right panel, Video S7), demonstrating that keratin filaments can be transported independently of vimentin.

**Figure 3.**
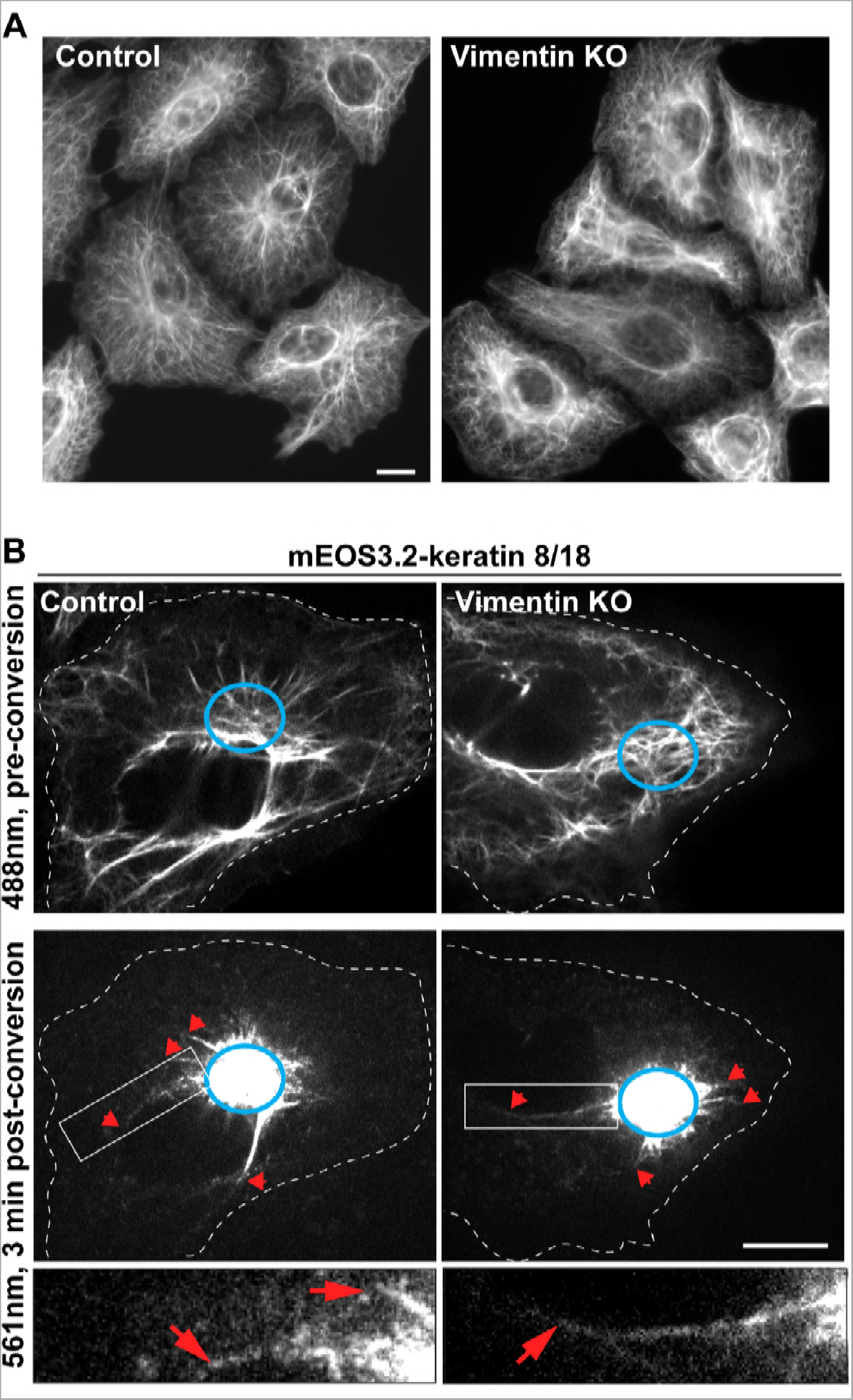
*Keratin filaments are transported in the absence of vimentin in A549 cells. A) Widefield microscopy imaging of keratin immunostaining in A549 cells (WT versus vimentin KO). B) Photoconversion experiment of mEos3.2-keratin 8/18 in A549 cells using spinning disk confocal microscopy. mEos3.2-keratin was photoconverted from green to red at the cell center (cyan circle). The top panels show keratin network in the green channel (488nm) before conversion and the bottom panels show the red channel (561nm) 3 minutes after photoconversion. Translocated photoconverted filaments are indicated by red arrows. The enlargemenst show that fully polymerized keratin filaments were transported even in absence of vimentin. Bars, 10µm.*

### Keratin and vimentin filaments are transported using the same mechanism

Our results have demonstrated that both vimentin and keratin filaments are transported along microtubules by kinesin-1. Typically, kinesin-1 binds to its cargo via kinesin light chains (KLC) (26-28). In a small number of cases, some cargoes bind to kinesin-1 tail and do not require KLC for transport (29, 30). To determine whether IFs utilize KLC as an adapter, we used genome editing to KO KLC1, the gene encoding the predominant isoform of KLC in RPE cells (Figure 4A). Immunostaining of vimentin and keratin filaments in RPE cells revealed that KLC1 KO did not affect the distribution of either type of IF network (Figure 4B, G). However, the human genome contains four genes encoding for KLC and their pattern of expression in RPE cells is not well established. Therefore, we decided to use an alternative approach, replacing the wild-type KIF5B in RPE cells with a truncated version of the motor lacking the region that recruits KLC to the kinesin-1 complex. To accomplish this, we deleted the heptad repeats (residues 775-802) of mKif5b responsible for KLC binding, creating (mKif5b^Δ775-802^-Emerald) (Figure 4C). Pull-down experiments and western blot analyses were performed to confirm that mKIF5B^Δ775-802^-Emerald did not interact with KLC. As described previously, KLC binding to kinesin-1 is required for their stability (31). As a consequence, neither KLC1 nor KLC2 were detectable by western blot analysis in crude extract of *KIF5B* KO cells (Figure 4D). We found that the rescue of *KIF5B* KO by expression of the full-length mKif5b-Emerald prevented degradation of KLC. In contrast, KLC 1 and 2 remained undetectable in lysates from cells expressing mKif5b^Δ775-802^-Emerald (Figure 4D). In addition to probing crude extracts, the kinesin-1 complex was enriched by pull down using GFP-binder and the pellets were probed for the presence of KLC1 and KLC2. These experiments showed that the full-length mKif5b-Emerald bound KLC1 and KLC2 while no KLC could be found even after enrichment of mKif5b^Δ775-802^-Emerald (Figure 4E).

**Figure 4.**
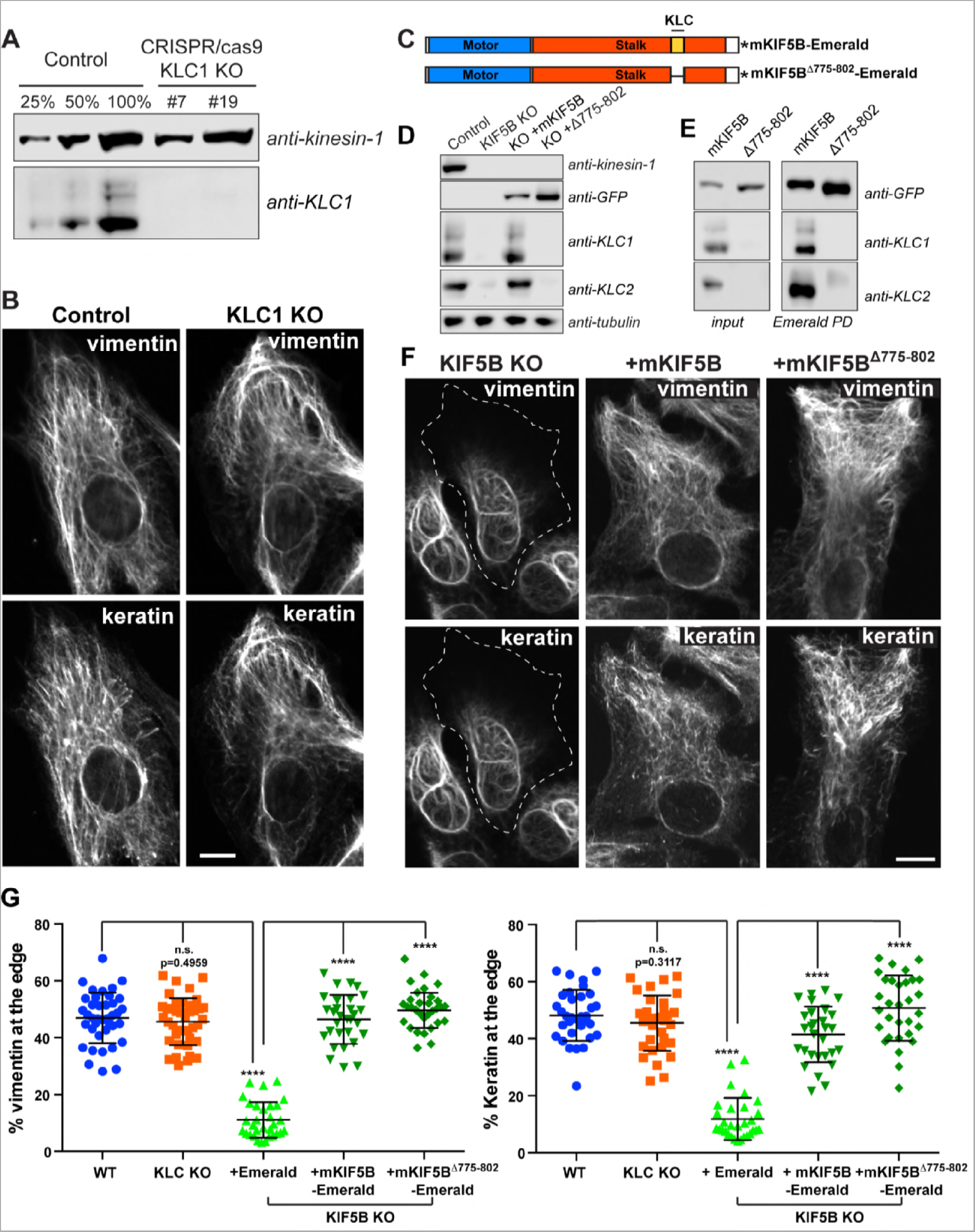
*Transport of vimentin and keratin filaments is independent of kinesin light chain. A) Western blot analyses using KLC-1 antibody showed the absence of KLC-1 in two different KLC-1 CRISPR clones (#7 and #19). KiF5 antibody was used as loading control. B) Confocal microscopy of vimentin and keratin immunostaining in RPE cells WT (control) and KLC-1 KO. C) Schematic representation of mKif5b-Emerald and mifF5b^Δ775-802^-Emerald. The domain in blue is the motor domain of mKif5b, the red part is the stalk which comprise the heptad repeat domain responsible for KLC binding in yellow. Note that this KLC binding domain is absent from the mKif5b^Δ775-802^-Emerald. The asterisk represents Emerald. D) Western blot analyses of kinesin-1, KLC1 and KLC2 showed that KLCs were absent from the lysate prepared from the KIF5B KO cells. Anti-tubulin was used as loading control. E) Endogenous KIF5B was replaced by mKif5b-Emerald or mKif5b^Δ775-802^-Emerald. The presence of KLC1 and KLC2 was determined by western blot of crude cell lysates (left panel) or after enrichment of the kinesin-1 complex by pull down using GFP-binder recognizing Emerald. F) Confocal microscopy imaging of vimentin and keratin immunostaining in RPE Kif5b KO cells after the expression of Emerald (KIF5B KO), mKIF5B-Emerald (+mKIF5B) or mKIF5B^Δ775-802^-Emerald (+Δ775-802). G) Graphs show the % of vimentin (left) or keratin (right) at the cell edge (Mean with SD; n >30 cells). Data are representative of at least two independent experiments. Statistical significance was determined using the Mann-Whitney test (****; p<0.0001)*

Immunostaining of vimentin and keratin IFs was employed to compare the efficiency of full-length mKif5b and mKif5b^Δ775-802^ in rescuing IF distribution. The images showed that removal of KLC and the region of kinesin tail that interacts with KLC had no effect on the capacity of mKif5b to rescue keratin or vimentin distribution (Figure 4F). This observation was reflected in the quantification of vimentin and keratin fluorescence intensity at the cell edge, confirming that both mKif5b and mKif5b^Δ775-802^ constructs rescued IF distribution to the same extent (Figure 4G). These results demonstrate that KLCs are not involved in the kinesin-dependent transport of IFs.

Since KLC was not required for IF transport, we concluded that a specific cargo-binding region of the kinesin tail might be involved. To determine which part of the KIF5B tail is important, we created three other truncated version of mKif5b fused to Emerald: mKif5b^1-773^, mKif5b^1-806^ and mKif5b^1-892^ (Figure 5A). All three constructs were properly expressed in *KIF5B* KO RPE cells and are likely functionally active as they accumulated at the cell periphery, where most of the microtubule plus-ends are located. This distribution was different from the distribution of full-length mKIF5B because the C-terminal truncations removed the autoinhibitory domain located at residues 937-952 of KIF5B (32), producing a constitutively active motor (Figure 5B).

**Figure 5.**
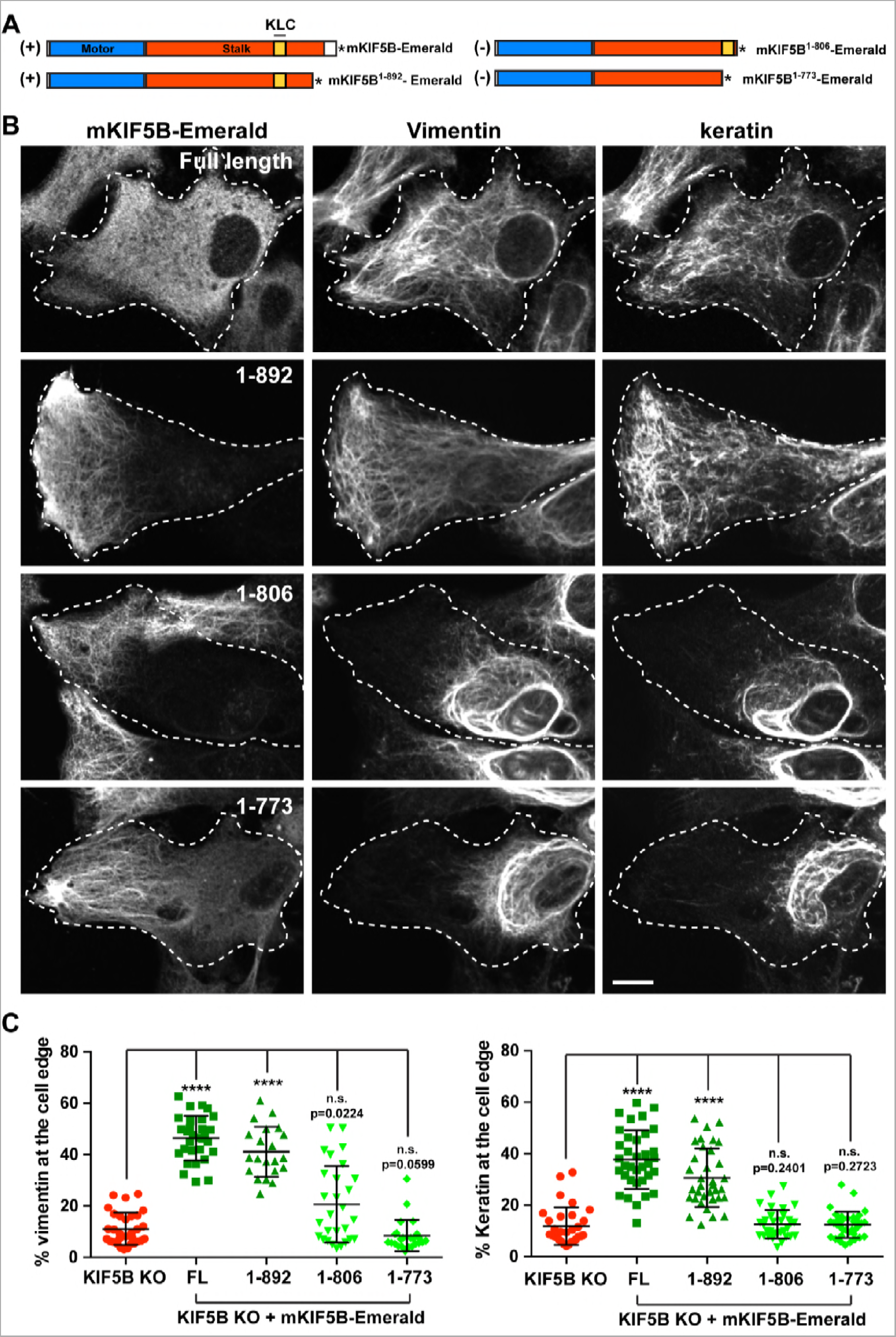
*Mouse Kif5b^1-982^ rescues IF distribution in KIF5B KO cells. A) Schematic representation of different truncations of the KIF5B tail. The mKIF5 constructs capable of rescuing vimentin and keratin distribution are marked by a (+). The asterisk represents Emerald. B) Confocal microscopy imaging of vimentin and keratin immunostaining in RPE KIF5B KO cells after the expression of mKif5b-Emerald (full length), mKif5b^1-892^-Emerald (1-892), mKif5b^1-806^-Emerald (1-806) or mKif5b^1-773^-Emerald (1-773). C) Graphs show the % of vimentin (left) or keratin (right) at the cell edge (Mean with SD, n >30 cells). Data are representative of at least two independent experiments. Statistical significance was determined using the Mann-Whitney test.*

The ability of these three truncations to rescue vimentin and keratin IF distribution in *KIF5B* KO cells was tested by immunostaining. Deletion of the last 70 residues of KIF5B, creating mKif5b^1-892^ did not prevent the ability of mKif5b to rescue the IF distribution, while more extensive truncations (mKif5b^1-773^, mKif5b^1-806^) inhibited the ability of mKif5b to properly position IF. These results and the rescue by mKIF5B^Δ775-802^, strongly support the presence of an IF binding between residues 803-892 of the KIF5B tail (Figure 5A). It is worth noting, that the constructs that rescued vimentin distribution were also able to rescue the distribution of keratin and vice versa.

## DISCUSSION

The development of fluorescent probes and advances in live cell imaging, have dramatically changed our understanding of IF dynamics. Traditionally considered as mostly static rigid structures, IFs are, in fact, highly dynamic, undergoing constant rearrangement by severing and re-annealing (7, 8), subunit exchange (33) as well as translocation of precursors and fully polymerized filaments (8, 12, 34, 35). In this paper, we established that kinesin-1 is essential to vimentin IFs transport along microtubules. In *KIF5B* KO cells, vimentin filaments are depleted from the cell edge (Figure 1A) and active transport of vimentin filaments from the cell center to the cell periphery is no longer observed (Figure 1C). Furthermore, we showed that even fully polymerized keratin IFs undergo constant transport along microtubules powered by kinesin-1 (Figure 2-3). Finally, we established, that KLC was not involved in vimentin or keratin IF transport by kinesin (Figure 4) and that the same region of the kinesin tail is required for both keratin and vimentin transport (Figure 5).

### Contribution of microtubules to keratin network dynamics

The dynamics of the keratin filament network has been extensively studied (25). It has been clearly demonstrated that actin dynamics are responsible for the retrograde transport of keratin filament precursors formed at the focal adhesion and long filaments to the perinuclear region (18, 20, 22, 36). Experiments using fluorescence recovery after photobleaching (FRAP) have been very powerful to decipher the role of subunit exchange during the cycle of assembly and disassembly of the entire keratin network (19, 37). This model was recently recapitulated *in vivo* in an elegant study using YFP-tagged keratin in murine embryos (38).

In our study, we used photoconversion as an alternative approach to follow a small subset of individual filaments at the cell center, the site where the filament network is especially dense. By using this technique, we were able to observe for the first time the anterograde transport of fully polymerized keratin filaments.

Our data complement very well the published keratin dynamics, demonstrating the contribution of microtubule-dependent transport from the cell center to the periphery. We believe that the contribution of microtubule-dependent transport to keratin IF dynamics is dependent on the physiological context. We looked at keratin dynamics in two different cell lines; RPE cells, that are highly motile, and A549 carcinomas cells, which are stationary and tend to form cell-cell contacts like typical epithelial cells. We observed keratin filament transport in both cell types, but it was more most robust in RPE cells, raising the possibility that microtubule-dependent transport of keratin filaments could be upregulated as the cells change shape. We speculate that anterograde transport of keratin filaments in epithelial cells delivers filaments to newly formed areas of cytoplasm in migrating epithelial cells.

### Kinesin-1 as a universal transporter of intermediate filaments

In this paper, we established using *KIF5B* KO that kinesin-1 is the anterograde motor responsible for the transport of vimentin and keratin IFs. A role for kinesin-1 has been suggested previously using less specific or less efficient approaches, such as antibody injection that inhibits multiple kinesins (9), or shRNA that causes incomplete knock down (13). In both cases, we could not exclude the possibility that kinesin-1 was cooperating with other anterograde motors to transport vimentin. Antibody injection caused a dramatic retraction of vimentin filaments that could be explained by the inhibition of multiple kinesins, and the incomplete retraction of vimentin filaments observed after *KIF5B* knock down could have been explained by the action of a second kinesin motor capable of moving vimentin filaments, or incomplete kinesin-1 depletion. In our current study, the KO of *KIF5B* completely removed kinesin-1 from RPE cells, resulting in the complete inhibition of not only vimentin, but also keratin transport along microtubules and the retraction of the two IF networks to the perinuclear region. This retraction is very likely caused by actin retrograde flow as demonstrated previously during nocodazole treatment (39), or by retrograde transport along microtubules (8, 40) and is normally counteracted by kinesin-driven anterograde transport.

Kinesin-1 KO has been demonstrated to have a major impact in at least two different tissues by affecting different types of IFs. Conditional KO of the neuronal isoform of kinesin-1, *Kif5a* in mouse neurons inhibits neurofilament transport into the axon leading to the accumulation of the three neurofilaments (NFs) NF-H, NF-M and NF-L in the cell body causing neurodegeneration and premature death of the animal (15, 41). More recently, conditional KO of *Kif5b* in mouse myogenic cells prevents the recruitment of desmin and nestin IFs to the growing tip of the myotube during muscle formation associated with severe skeletal muscle abnormalities and heart failure (16). Based on our study and others, we can conclude that all types of IFs, except perhaps the nuclear lamins, are kinesin-1 cargoes.

### Are all IFs transported along microtubules using the same mechanism?

Using mKIF5B truncations to rescue vimentin and keratin IFs distribution in *KIF5B* KO cells, we determined that KLC was not involved in the transport of these two types of IFs and that a region between residues 803-892 of the kinesin tail was required (Figure 4-5). These findings corroborate a previous IF study showing that proper localization of desmin- and nestin-containing IFs in the myofibril is independent of KLC and can be rescued with a truncated version of KIF5B lacking the last 73 residues of the kinesin tail (1-890)(16). This suggests that several types of IF probably bind to the same region of the kinesin tail, reinforcing the hypothesis that all IFs are transported along microtubules by a common mechanism. However, it is unknown whether IFs bind to kinesin-1 directly, or if this binding is mediated by an adaptor protein. Direct interaction between the IF protein desmin and the kinesin tail has been demonstrated *in vitro* (16), suggesting that it could be the case for other types of IF. If an adaptor is involved, it likely links all types of IF to the region 803-892 of the kinesin tail. Whether IF binding to kinesin-1 is direct or adaptor-mediated, the region of the IF proteins responsible for the interaction with kinesin-1 needs to be identified. Since all IF types are potential kinesin-1 cargoes, it is plausible that the recognition domain for kinesin-1 is located in the central α-helical rod domain that is highly conserved in IF proteins (42).

There is compelling evidence that IFs functions go well beyond the control of mechanical integrity, as they are becoming key players in the signal transduction of stress-related and other cellular responses (43). In that context, the proper delivery of IFs to sub-cellular locations might be a key requirement for their ever-growing list of functions.

## MATERIALS AND METHODS

### DNA constructs

mEos3.2-vimentin in pQCXIN was described before (8). To create mEos3.2-keratin 8 and mEos3.2-keratin 18, Eos3.2 was amplified from mEos3.2-vimentin by PCR with Phusion polymerase (Clontech) and joined to pQCXIN cut with NotI by InFusion (Takara). The resulting vector was cut with BamHI. The keratins were amplified from pcDNA-keratin 8 or 18 by PCR with Phusion polymerase and joined to the BamHI cut pQCXIN with InFusion.

Mouse KIF5B (mKIF5B) cDNA was provided by Addgene (pKIN1B plasmid #31604). To create mKIF5B deletion, mKIF5B was amplified by PCR using the forward primer ATAAGAATGCGGCCGCTTCCAGAAAGATGGC together with one of the following reverse primers. For mKIF5B full length, ATGGATCCCACGACTGCTTGC CTCCACCAC; mKIF5B^1-773^, ATGGATCCCAGTCTTGCATAACCGTGAGCT; mKIF5B^1-806^, ATGGATCCCAGTCCTGAACAAAGAGCTTAC; mKIF5B^1-892^, ATGGATCCCAACGGTCT CGAGATGCATTTT. For mKIF5B^Δ775-802^, the complementary oligos CACGGTTATG CAAGACAGATTTGTTCAGGACTTGCCTACCAGGGTGAAAAAGAGCGCCGAGG and TCGACCTCGGCGCTCTTTTTCACCCTGGTAGCCAAGTCCTGAACAAATCTGTCTTGCATAACCGTGAGCT were phosphorylated, annealed and inserted into the SacI/SalI restricted sites of the mKIF5B cDNA. The deletion mutant was amplified by PCR using the same primers as mKIF5B full length. All the Emerald tagged mKIF5B constructs were generated by replacing vimentin from vimentin-Emerald pQCXIN (8) by the mKIF5 constructs using the AgeI/BamHI sites.

To generate untagged mouse vimentin (mVimentin) in pQCXIP, mVimentin was amplified using the primers CGCACCGGTATGTCCACCAGGTCCGTG and CGGAATTCCTTATTCAAGGTCATCG. The PCR fragment was digested and inserted into the AgeI/EcoRI sites of the pQCXIP vector. The generation of the Y117L-vimentin mutant cDNA has been described (44).

### Antibodies and reagent

Chicken polyclonal anti-vimentin (PCK-594P) is from BioLegend (Dedham, MA); mouse pancytokeratin (C2931) is from Sigma (St. Louis, MO); rabbit polyclonal KLC1 (19028-1-AP) and KLC2 (17668-1-AP) antibodies are from Proteintech (Rosemont, IL). CT and HD polyclonal antibodies against kinesin are a kind gift from Fatima Gyoeva (Institute of Protein Research, Russian Academy of Sciences, Moscow, Russia). Rhodamine-conjugated phalloidin, MitoTracker Deep Red and LysoTracker Deep red dyes are from Invitrogen Molecular Probes (Eugene, OR)

### Cell lines and CRISPR/Cas9 knock out

All cells were maintained at 37°C in 5% CO2. RPE cells were maintained in DMEM supplemented with 1 mM sodium pyruvate (Gibco) and 10% fetal bovine serum (FBS, Neuromics); Human lung carcinoma A549 WT and Vimentin KO cells were generously provided by Dr. Karen Ridge (Northwestern University, Chicago, IL). A549 cells were maintained in DMEM (Corning Cellgro, Mediatech Inc.) supplemented with 10% FBS and 10mM HEPES (Gibco).

All vectors for CRISPR/cas9 genome editing are from GenScript (Piscataway, NJ). The Kif5b KO and klc1 KO cell lines were created by the transduction of RPE or A549 cells with lentivirus carrying LentiCRISPR V2 KIF5B gRNA1 (target sequence CTATACCTTGTGCTCGAAGC) or LentiCRISPR V2 KLC1 gRNA1 (target sequence GAAGCAGAAACTGCGTGCGC). Lentivirus were produced in HEK 293 FT cells transfected with the LentiCRISPR V2 vector containing cas9 and KIF5B or KLC1 gRNA together with the helper plasmids pVSVG (Clontech) and pPAX2 (Imgenex) encoding the gag/pol and env proteins required for virus production. Virus-containing medium was collected and filtered 48 hours after the transfection of the packaging cells. 8µg/mL of polybrene (Sigma) was added to the freshly collected viruses and RPE or A549 cells were incubated for 6 hours with this virus/medium/polybrene mixture. Two days later, infected cells were selected using 5µg/mL puromycin for 1 week and survivor cells were plated at 1 cell/well.

To KO vimentin in RPE, cells were transfected with pGS-vimentin gRNA1-Neo (target sequence ATTGCTGACGTACGTCACGC) using Lipofectamine 3000 according to the manufacturer’s instructions. 48h after transfection, transfected cells were selected using 1mg/mL G418 for two weeks and survivor cells were plated at 1cell/well. For all CRISPR cell lines, single colonies were amplified, lysed and tested for knock out by western blot.

mEos3.2-vimentin and mEos3.2-keratin 8/18 were stably expressed in RPE (WT and *KIF5B* KO) or in A549 by retroviral transduction. Retroviruses were produced as described above except that the helper plasmids pVSVG (Clontech) and pCL-Eco (Imgenex, San Diego, CA) were used instead. Transduced cells were selected using 2mg/ml G418 for 1 week. Retrovirus transduction of RPE KIF5b KO was performed to replace WT KIF5b by the different mouse KIF5b constructs listed above. mVimentin and mVimentin^Y117L^ were also expressed in RPE vimentin KO cells by retrovirus transduction.

### Enrichment of the kinesin-1 complex by pull down using GFP-binder

*KIF5B* KO cells expressing mKif5b-Emerald of mKif5b^Δ775-802^-Emerald from two sub-confluent 100mm dishes lysed in 1mL of ice-cold RIPA buffer (50mM Tris pH 7.4, 150mM NaCl, 1% Triton, 0.5% Na Deoxycholate, 0.1% SDS, 10mM NaPPi, 1.5mM NaVO_3_, 1mM PMSF) supplemented with peptidase inhibitors (CLP, chymostatin, Leupeptin and Pepstatin A 20µg/mL). The cell lysates were centrifuge at 20,000g for 5 min. The soluble fraction was incubated for 4 hrs at 4°C with 30µL sepharose beads conjugated with single chain GFP antibody (GFP-binder, GFP-Trap-M from Chromotek). Beads were washed 3 times with RIPA-base (50mM Tris pH 7.4, 150mM NaCl, 1% Triton X-100, 10mM NaPPi, 1.5mM NaVO_3_, 1mM PMSF) supplemented with CLP. The kinesin-1 complex was pulled down via binding of Emerald from mKIF5B-Emerald to the GFP binder beads by centrifugation at 3000xg and resuspended in 30µL of Laemmli buffer (5% SDS, 0.1mM Tris pH 6.8, 140mM β-mercaptoethanol, 0.25% glycerol). Samples were boiled for 5 minutes and analysed by western blot.

### Immunostaining for widefield and confocal microscopy

Cells were plated on glass coverslips to desired confluence ∼16hr before fixation. For vimentin and actin co-staining plus keratin and actin co-staining, cells were fixed with 3.7% formaldehyde in CSK buffer (100mM NaCl, 300mM Sucrose, 3mM MgCl2, 10mM Pipes pH 6.8) supplemented with 0.1% Triton X-100 for 10 min. For vimentin and keratin co-staining, cells were fixed with ice-cold MeOH for 5 min at −20°C. Fixed cells were then further extracted in 0.2% Triton X-100 in PBS before staining. Immunostaining was performed in wash buffer (TBS supplemented with 1% BSA, 0.1% Triton-X100) as described previously (12).

Images of fixed cells were captured on a Nikon Eclipse U2000 inverted microscope (Nikon Instruments, Melville, NY, USA) equipped with Ph3 40x/1.0 NA objective and a CoolSnap ES CCD camera (Roper Scientific, Planegg/Martinsried, Germany), driven by Nikon Elements software. Fluorescence excitation was achieved using a mercury lamp. Images of vimentin and keratin co-staining were collected from a Zeiss confocal LSM510 META (Carl Zeiss, Jena, Germany) microscope with oil immersion objective lenses (Plan-Apochromat, 60×, 1.40 numerical aperture NA; Carl Zeiss).

### Vimentin and keratin distribution measurement

Widefield microscopy images of vimentin or keratin and actin co-staining captured as described above were analysed using the FIJI software version 2.0. For at least 30 cells per condition, a single line, three pixels in width, was traced manually from the center of the nucleus to the cell edge as delineated by the F-actin staining (See Figure S1). The intensity values along this line were obtained using the “plot profile” plugin from FIJI. The mean fluorescence intensity and the positions along the line for each cell were normalized from 0 to 1 using Matlab. The normalized data were transferred to Prism 7 software (GraphPad Prism, GraphPad Software, San Diego CA) for further analyses. The percentage of signal at the edge was calculated by dividing the area under the curve (AUC) between positions 0.5 to 1 by the total AUC between positions 0 to 1. Data are representative of two or more independent experiments and are shown as the mean with SD (n>30). Statistical significance was determined using the non-parametric Mann-Whitney test with a confidence interval of 95%. This test analysis compares the distributions of two unpaired groups.

### Keratin network spreading measurement

Keratin network spreading analyses were performed on confocal images of vimentin and keratin co-staining, acquired with 100x objective lens as described above using the FIJI software version 2.0. The vimentin signal was used to manually trace the area of single cell using the polygon selection tool of FIJI. High contrasted image of the non-specific staining was used to determine to outline of the vimentin KO cells. The corresponding channel for the keratin signal was auto-threshold using the Li method to determine the area (in pixel) covered by keratin signal. The percentage of keratin spreading was determined by dividing the keratin area by the total area of the cell. Data are representative of at least two independent experiments and are shown as the mean with SD (n>30). Statistical significance was determined using the non-parametric Mann-Whitney test with a confidence interval of 95%. This test analysis compares the distributions of two unpaired groups.

### Live cell imaging and photoconversion

For all live-cell experiments, cells were plated on glass coverslips ∼16hr before imaging. Cells were maintained at 37˚C in 5% CO_2_ during imaging using a Tokai-Hit stage-top incubator (Tokai-Hit, Fujinomiya City, Japan) and Okolab gas mixer (Okolab, Naples, Italy).

Confocal images were collected on a Nikon Eclipse U2000 inverted microscope equipped with a Yokogawa CSU10 spinning-disk confocal head (Yokogawa Electric Corporation, Sugar Land, TX), a Plan Apo 100×/1.45 NA objective, an Agilent MLC 400 laser set (including 488nm and 561nm lasers; Agilent Technologies, Wood Dale, IL), 89 North Heliophor pumped phosphor light engine at 405nm (Chroma Technology, Bellows Falls, VT) to drive photoconversion, and an Evolve EMCCD (Photometrics, Tucson, AZ) driven by Nikon Elements software. Photoconversion mEos3.2-keratin 8/18 from green to red was performed using illumination from a Heliophor LED light source in the epifluorescence pathway filtered with a 400-nm filter and confined by a diaphragm. Photoconversion time was 5 s and the zone was 10 μm in diameter, which was positioned at the cell center. Time-lapse sequences were acquired at 15s intervals for 3 min using 561nm laser. Images were analyzed in Fiji, and assembled in Illustrator.

Imaging of mEos3.2-vimentin was performed using TIRF on a Nikon Eclipse U2000 inverted microscope equipped with a Plan-Apo TIRF 100x 1.49 NA objective and a Hamamatsu CMOS Orca Flash 4.0 camera (Hamamatsu Photonics K.K. Hamamatsu City, Japan), controlled by NIS-Elements AR 4.51.01 software (Nikon, Melville, NY, USA). The angle of a 561 nm laser was manually adjusted until near total internal reflection was reached as judged by imaging of photoconverted mEos3.2-vimentin expressing cells. For photoconvertion, cells were exposed for 3 sec to UV light from a mercury arc in the epifluorescent light path filtered though a 400nm excitation filtered and spatially restricted by a pinhole in the field diaphragm position. Time-lapse sequences were acquired at 15 sec intervals for 3 min using the 561nm laser.

For each photoconversion experiments, at least ten cells were photoconverted and each condition was repeated in at least three independent experiments. The representative photoconversion data are shown for each condition.

## ACKNOLEDGEMENT

Research reported in this publication was supported by the National Institute of General Medical Science of the National Institutes of Health under awards P01GM09697 and R01 GM52111. The authors would like to thank Dr. Karen Ridge (Northwestern University, Chicago, IL) for the A549 WT and vimentin KO cells, Fatima Gyoeva (Institute of Protein Research, Russian Academy of Sciences, Moscow, Russia) for the CT and HD polyclonal antibodies against kinesin and David Kirchenbuechler from the Center for Advanced Microscopy/Nikon Imaging Center (Northwestern University, Chicago, IL) for help with quantification of IF distribution.

## Supplemental Material

**Figure S1.**
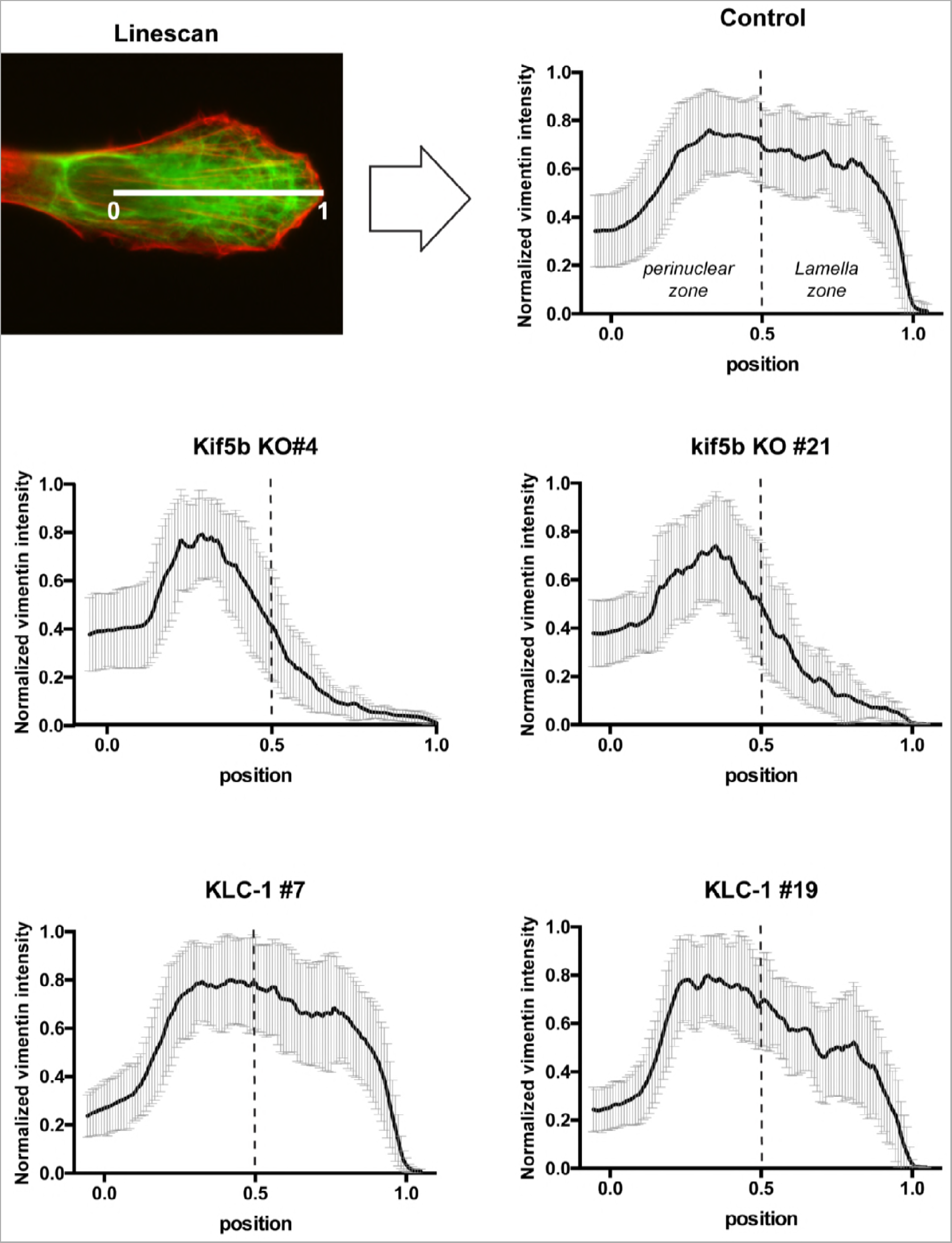
***Plot profile analysis of filament distribution**. The illustration show an example of two-color image used for the quantification. The red channel (F-actin labeling) was used to trace a 3 pixels-wide line from position 0 (middle of the nucleus) and 1 (cell edge). The intensity profile fro the green channel (vimentin staining) along that line was obtained using the plot profile plugin in FIJI. The results were normalized using MatLab and each graph represents the normalized data from at least 30 cells per condition. The dashed line represents the midpoint of the line that was used to separate the element present at the cell edge (between 0.5 to 1) from the element in the perinuclear region (between 0 and 0.5). These graph were used to calculate the area under the curve at the cell edge versus total presented in figure 1. The same method was used to quantify the distribution of keratin (Figure 2) and mitochondria (Figure S2).*

**Figure S2.**
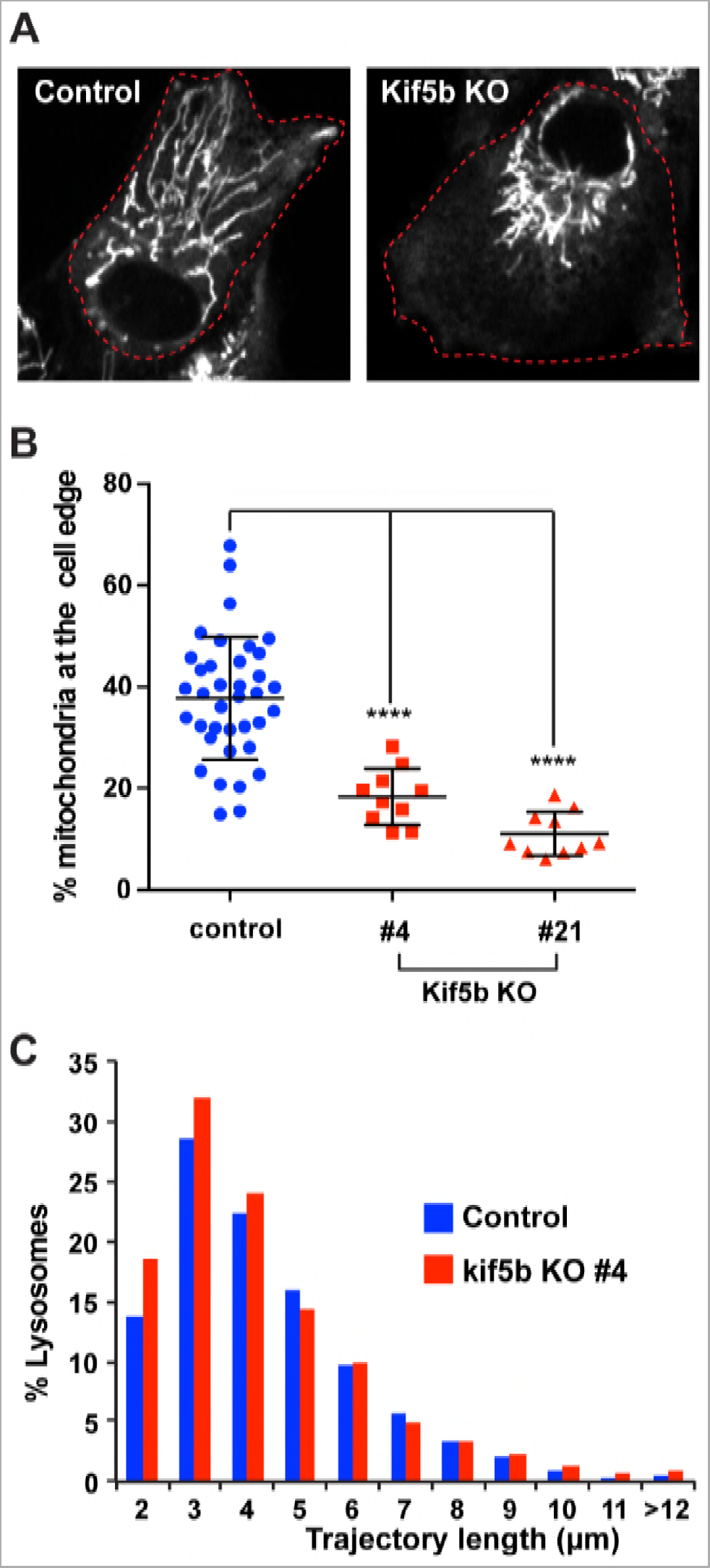
***KIF5B KO affects mitochondria distribution but not lysosome transport**. A) RPE WT and KIF5B KO were incubated with MitoTracker Deep red (1:20 000 for 10 minutes) and observed by confocal microscopy. The red dashed lines delineate the cell periphery. B) Mitochondria distribution was quantified as described for vimentin and keratin except that phase contrast images were used to determine the cell edges. The graph shows the depletion of mitochondria from the cell edge in two different KIF5B KO clones (#4 and #21). Statistical significance was determined using Mann-Whitney test (****; p<0.0001). C) RPE WT and KIF5B KO were incubated with LysoTracker Deep red (1:50 000 for 10 minutes) and lysosomes were imaged by confocal microscopy every second for 1 minutes. The lysosome trajectory length was measured in 11 cells per condition (n>2500 lysosomes).*

**Figure S3.**
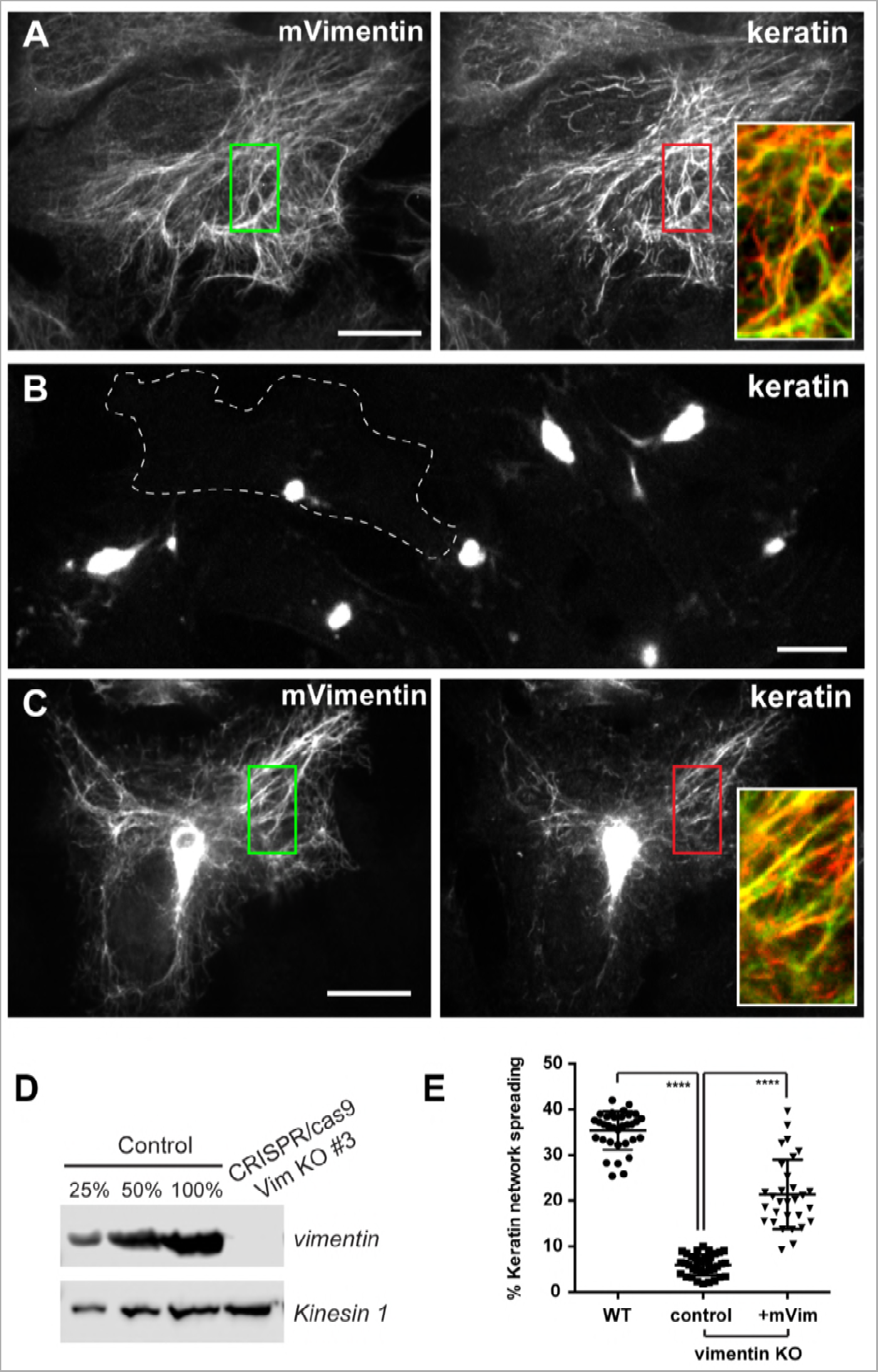
***Keratin filaments collapse in absence of vimentin in RPE cells**. A-C) Confocal images of keratin and vimentin immunostaining in RPE WT (A), vimentin KO (B) or vimentin KO expressing mVimentin (C). The boundary of on cell in (B) was delineated with a dashed line to highlight the intensity of the keratin filaments collapse. Enlargements of the inset emphasize the connection between the two networks. Bars, 10μm. D) Western blot analyses using vimentin antibody shows the absence of vimentin in clone #3 after vimentin KO using CRISPR/cas9. Kinesin-1 is used as loading control. E) The percentage of keratin network spreading represent the fraction of the keratin staining coverage for each cell for at least 30 cells per conditions. Statistical significance was determined using the Mann-Whitney test (****; p<0.0001).*

### Video S1.

Vimentin filaments are transported in RPE cells. A 10 µm-diameter area of a mEos3.2-vimentin expressing cell was photoconverted from green to red with 405nm light (see cyan circle). The movie shows the transport of red vimentin filaments outside of the photoconverted area every 20 sec after photoconversion for 3 min.

### Video S2.

KIF5B KO inhibites vimentin filaments transport. A 10 µm-diameter area of a *KIF5B* KO RPE cells expressing mEos3.2-vimentin was photoconverted from green to red with 405nm light (see cyan circle). Pictures of the red channel were taken every 20 minutes for 3 min. The movie shows that vimentin filaments are confined inside the photoconverted area when KIF5B is absent.

### Video S3.

Keratin filaments are transported in RPE cells. A 10 µm-diameter area of a mEos3.2-keratin expressing cell was photoconverted from green to red with 405nm light (see cyan circle). The movie shows the transport of red vimentin filaments outside of the photoconverted area every 15 sec after photoconversion for 3 min.

### Video S4.

Keratin filament transport requires microtubules. mEos3.2-keratin expressing cells were treated with 10µM nocodazole for 3h to depolymerize microtubules. A 10 µm-diameter area of a nocodazole-treated cell was photoconverted from green to red with 405nm light (see cyan circle). Pictures of the red channel were taken every 20 minutes for 3 min. The movie shows that keratin filaments are confined inside the photoconverted area in absence of microtubules.

### Video S5.

KIF5B KO inhibites keratin filaments transport. A 10 µm-diameter area of a *KIF5B* KO RPE cells expressing mEos3.2-keratin was photoconverted from green to red with 405nm light (see cyan circle). Pictures of the red channel were taken every 20 minutes for 3 min. The movie shows that keratin filaments are confined inside the photoconverted area when KIF5B is absent.

### Video S6.

Keratin filaments are transported in A549 cells. A 10 µm-diameter area of a mEos3.2-keratin expressing cell was photoconverted from green to red with 405nm light (see cyan circle). Pictures of the red channel were taken every 20 sec after photoconversion for 3 min. The red arrows point some red keratin filaments that are being transported outside of the photoconverted area.

### Video S7.

Keratin filaments are transported in absence of vimentin in A549 cells. A 10 µm-diameter area of a vimentin KO A549 cells expressing mEos3.2-keratin was photoconverted from green to red with 405nm light (see cyan circle). Pictures of the red channel were taken every 20 minutes for 3 min. The red arrows point some red keratin filaments that are being transported outside of the photoconverted area even in cells lacking vimentin.

